# Identification of Casiopeina II-gly secondary targets through a systems pharmacology approach

**DOI:** 10.1101/327718

**Authors:** Guillermo de Anda-Jáuregui, Jesús Espinal-Enríquez, Junguk Hur, Sergio Antonio Alcalá-Corona, Lena Ruiz-Azuara, Enrique Hernández-Lemus

## Abstract

Casiopeinas are a group of copper-based compounds designed to be used as less toxic, more efficient chemotherapeutic agents. In this study, we analyzed the *in vitro* effects of Casiopeina II-gly on the expression of canonical biological pathways. Using microarray data from HeLa cell lines treated with Casiopeina II-gly, we identified biological pathways that are perturbed after treatment. We present a novel approach integrating pathway analysis and network theory: The Pathway Crosstalk Network. We constructed a network with deregulated pathways, featuring links between those pathways that crosstalk with each other. We identified modules grouping deregulated pathways that are functionally related. Through this approach, we were able to identify three features of Casiopeina treatment: a) Perturbation of signaling pathways, related to induction of apoptosis; b) perturbation of metabolic pathways, and c) activation of immune responses. These findings can be useful to drive new experimental exploration on their role in adverse effects and efficacy of Casiopeinas.

## Introduction

The need for new and better pharmacological therapies for the treatment of cancer still exists. While there are succesful specific treatment frameworks against some types of cancer, such as anti-hormonal therapy in breast cancer [14], or monoclonal antibodies targeting aberrant receptors [62], the intrinsic heterogeneity found in cancer forces the use of highly toxic chemotherapeutic regimes. One of the main goals in precision medicine is to focus in minimizing cancer treatment’s side effects [1]. Indeed, the wide range of adverse effects resulting from cancer therapy have an impact on therapeutic adherence [21, 26] and general quality of life for patients and their families.

Casiopeinas are a group of copper-based chemical compounds with cytotoxic activity. Casiopeinas were developed based on the rationale that copper compounds, unlike other metallic-based therapies, are more readily metabolized [15]; this property decreases the incidence of side effects found in several other chemotherapies [33]. These compounds have demonstrated efficacy as viable alternatives for the treatment of cancer, including lung [29] and cervix [54]. Studies have also found that these compounds have a mechanism of action that involves the induction of apoptosis through DNA damage, activation of reactive oxygen species, and mitochondrial dysfunction [29, 54, 43, 7, 4].

From a medicinal chemistry point of view, the use of genomics as a tool for the description of compounds is useful if it adds a new layer of information about the activity of these compounds. Based on this premise, in previous works [54, 17] the effects of Casiopeina II-gly (Cas II-gly) on gene expression have been measured on HeLa cell lines treated with this compound. Gene expression profiling identified activation of apoptotic pathways and inhibi-tion of cell proliferation processes. However, these previous works did not take into account that pathways are not isolated, but rather they interact through the phenomenon of pathway crosstalk.

In the present study, we further analyzed the aforementioned HeLa dataset, using a novel integrative approach that combines pathway analysis and network theory: the *Pathway Crosstalk Network* (***PXN***). This PXN contextu-alizes the processes and functions altered by Cas II-gly treatment *in vitro*, and the interactions taking place between them.

With this approach, we identified unseen alterations that may point to secondary mechanisms and targets of Casiopeinas, either as potential origins of adverse effects, or as secondary sites that contribute to a therapeutic effect and may offer synergistic opportunities. In particular, we identified the following systemic effects of Cas II-gly treatment: a) perturbation of signaling pathways related to apoptosis; b) perturbation of metabolic pathways, and c) activation of immune responses.

## Materials and Methods

### Dataset

The microarray rawdata was obtained from the NCBI Gene Expression Omnibus (Accession ID: GSE41827) [54]. In the original study, HeLa cell cultures were treated with 40 *μ*M of Cas II-gly (the previously identified IC_50_), and incubated for 6 hours before RNA extraction. RNA transcriptomes were quantified following the standard procedure for GPL570 platform (Affymetrix HGU-133_Plus_2). Experiments were performed in triplicate for both treated and untreated HeLa cell cultures. Microarrays were re-analyzed following a standard bioinformatics pipeline consisting of summarization and normalization using fRMA [35].

### Pathway deregulation analysis

Our main interest was the effect of Cas II-gly in gene expression at a functional level. With this in mind, we performed a pathway enrichment analysis strategy. We chose to use the KEGG database [30] as the source of our path-way collection; these were retrieved from the Molecular Signature Database [51]. We used the Generally Applicable Geneset Enrichment (GAGE) algorithm [32] to identify perturbations after Cas II-gly treatment. Unlike other enrichment strategies, GAGE performs comparisons considering the expression levels of all genes experimentally meassured, reducing biases that may be induced by the use of thresholds.

Importantly, GAGE allows the detection of enrichment either if there is an overall change of expression in the pathway molecules in a similar direction (over or under expressed) as well as in the case where molecules in the pathway may show changes in different directions. For this work we consider deregulated pathways in all directions, namely, upregulated, downregulated, or mixed. False discovery rate (FDR) q-value < 0.1 was used as the statistical significance cutoff.

### Pathway crosstalk network construction and community detection

Based on the idea that biological pathways are not isolated, but in fact communicate through molecular interactions, as well as that many molecules participate in different pathways, we propose the use of a *pathway crosstalk network* (***PXN***) as a tool for the exploration of Cas II-gly effects.

PXN is a graph composed of nodes representing pathways (in this case, pathways as defined in the KEGG database); these pathways were identified as perturbed by GAGE. Links in this network are drawn between pathways if they crosstalk: that is, if they share any molecule. The weight of links represents the similarity between pathways (molecules shared), measured using the Jaccard index (the ratio between the size of the union over the size of the intersection of two sets); a larger link weight suggests more crosstalk events between two pathways [14].

The structure of a PXN corresponds to a non trivial, in terms of the number of nodes and links and their distribution, as it is derived from the biological relationships between pathways. Therefore, the topological properties of a PXN are helpful to identify different roles of pathways in a specific biological context, in this case, the treatment with Cas II-gly. There are two global properties particularly relevant for this study: the number of connected components, i.e., islands, and the characteristic path length [57], which indicates the average minimum number of steps required to reach a node in the network from any other node. Together these features measure the information flow between pathways and the capacity of a given pathway to influence others.

Global organization patterns in networks involve the emergence of structural sub-units called *communities*: subsets of densely interconnected nodes [20, 41, 19, 3]. For a PXN, a community represents a set of functionally-related processes. For their detection we used the Infomap algorithm [45], since it is one of the most accurate and fastest algorithms according to the widely used Lancichinetti, Fortunato and Radicci (LFR) benchmark [31].

## Results and Discussion

We identified 78 pathways annotated in the KEGG database whose genetic expression was significantly altered after Casiopeina II-gly treatment. Of these, 41 pathways showed upregulation, 20 showed downregulation, and 17 showed a mixed effect in their overall expression pattern. We then mapped these pathways to a network representation as shown in Figure 1. The characteristics of this network is summarized in Table 1.

**Table 1:** Network of Pathways Perturbed by Casiopeina II-gly.

|  |  |
| --- | --- |
| Total Pathways | 78 |
| Upregulated | 41 |
| Downreagulated | 20 |
| Mixed perturbation | 17 |
| Edges | 712 |
| Connected Components | 1 |
| Isolated pathways | 2 |
| Char. Path Length ($\langle l \rangle$) | 2 |
| Communities | 9 |

**Figure 1:**
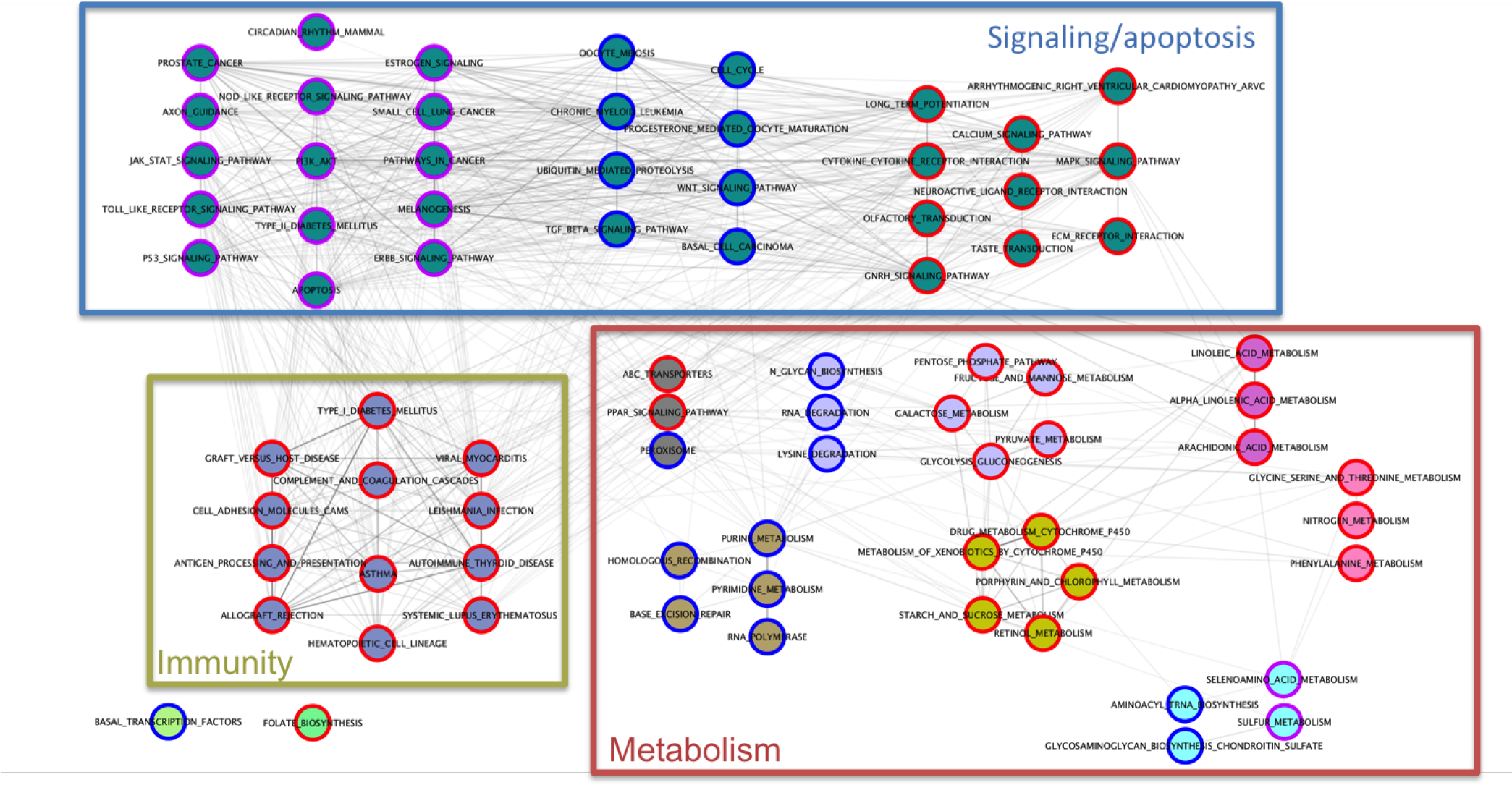
Network of Casiopeina II-gly perturbed pathways. Each node in the graph is a pathway. The border color represents the type of perturbation: red for upregulation, blue for downregulation, and purple for mixed effects. Community structures in the network were identified using the Infomap algorithm, represented by node color. Each frame shows the main activity of those pathways: Blue frame represents signaling pathways and apoptosis processes, bottom left yellow frame shows immunty-related pathways, the red frame represents metabolic pathways.

In this network we see the actions of Cas II-gly at a functional level. Even better, we observe such actions not as isolated features, but rather as an intricate network of interrelated processes. It is possible to identify processes that are functionally related grouped together. 76 pathways form the giant connected component and are organized in 9 communities reflecting common functionality. Furthermore, the network has a very short characteristic path length (< *l* >= 2). This suggests a fast and robust communication between different biological functions affected by Cas II-gly treatment. Interestingly, only two pathways affected by Cas II-gly treatment are disconnected from the rest of the network (bottom left in Fig. 1). In what follows, we will discuss the effects of Cas II-gly on these pathways and their possible pharmacological implications.

### The role of signaling pathways in the apoptotic activity of Casiopeina II-gly

The largest deregulated pathway community contains 33 pathways (top of Fig. 1), including signaling pathways and the apoptosis pathway. Induction of apoptosis at the cellular level is a well-known effect and mechanism of action for Casiopeina II-gly and other related compounds [15, 53, 36, 11, 52]. Indeed, the previous analysis of these microarray samples [54] identified apoptosis induction as a key process induced in the HeLa cells under Casiopeina II-gly treatment. Our analysis complements these previous findings, showing that a generalized perturbation of the molecules is involved in apoptosis by the aforementioned chemotherapy. We categorize this alteration as a mixed perturbation, which is congruent with the fact that apoptosis activation is not only manifested by an overexpression of apoptotic molecules, but also requires an underexpression of antiapoptotic molecules.

Twenty eight pathways in this community are directly connected to the apoptosis pathway. Interestingly, most of these are signaling pathways. Casiopeina II-gly has an overall activating effect on the MAPK signaling pathway [54]. Importantly, this pathway has the most connections to other pathways in our network (43), which is not surprising, as this signaling transduction system is widely known to be involved in a variety of responses [60]; activation of the MAPK pathway has been implicated in the cytotoxic activity of cisplatin[50]. In addition to the MAPK pathway, the calcium signaling pathway is also activated. The importance of intracellular calcium concentrations in apoptosis is well known [34, 37, 22, 16], and is consistent with Casiopeina activity. Alterations in calcium signaling could also be related to the activation of the disease pathway, arryhtmogenic right ventricular cardiomyopathy, whose activation could be a flag for a potential cardiotoxic activity of the compound.

We found downregulation in the Wnt and TGF-beta signaling pathways. The Wnt pathway regulates apoptosis, with Wnt-1 being an antiapoptotic signal [12, 24, 38]. Its inhibition is then consistent with Casiopeina inducing apoptosis. TGF-beta is known to play different roles on apoptosis and cell survival depending on the cellular context [27, 58, 42, 28, 59]. Casiopeina’s effect on this pathway may be reflecting an action upon its proapoptotic function. More importantly, there are reports of the antitumorigenic effects of TGF-beta inhibitors [25], opening the possibility for Casiopeina II-gly to have therapeutic effect by also targeting this pathway.

We also identified mixed effects on the expression of the PI3K-Akt, JAK-stat, NOD-like receptor, ERBB receptor, estrogen receptor, and P53 signaling pathways: all with established roles in tumor development. Overall, we have shown Casiopeina II-gly can affect a variety of signaling pathways which may be directly related to their pharmacological effects. A simplified diagram of these relationships can be seen in Figure 2.

**Figure 2:**
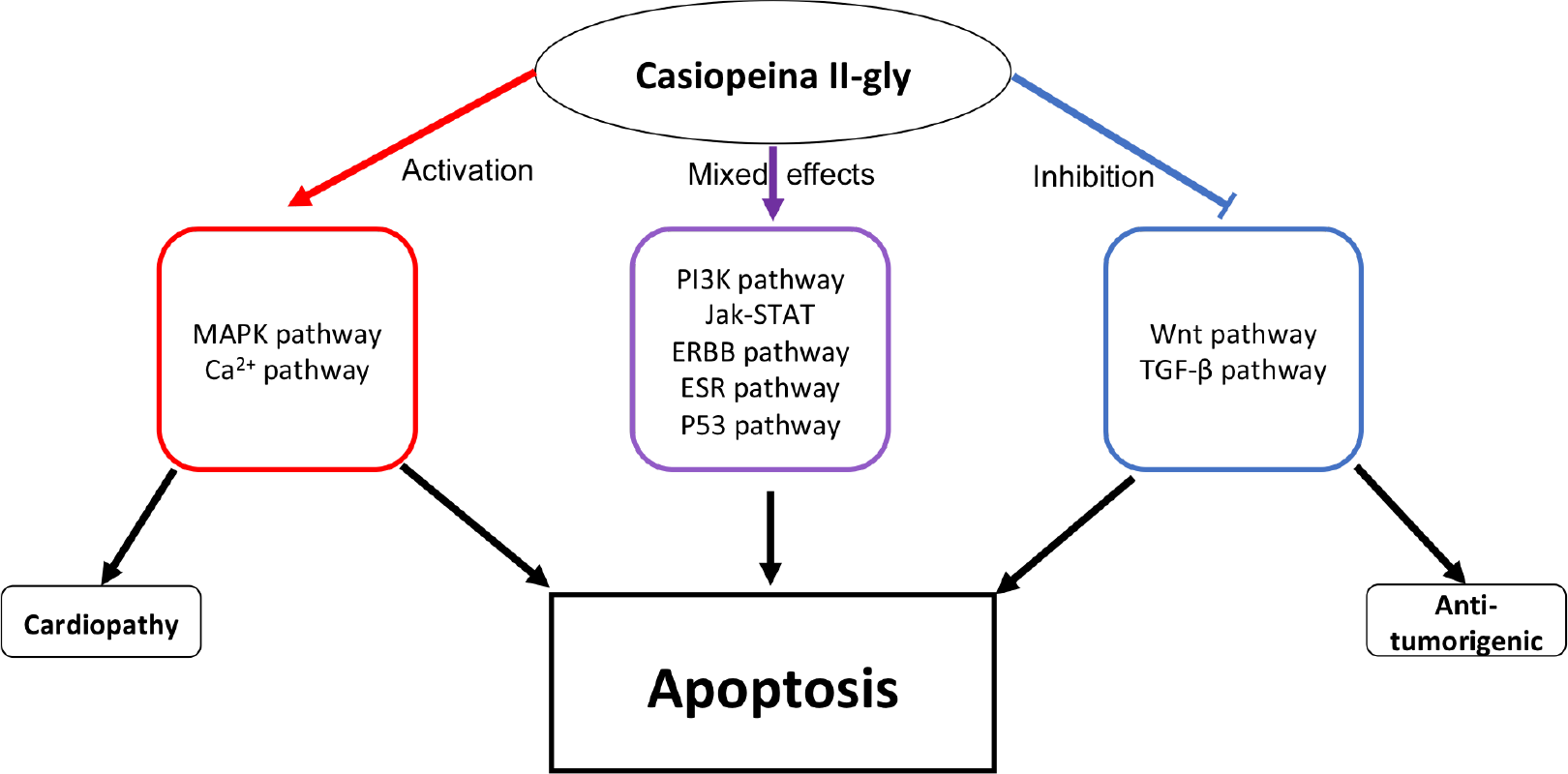
Effects of Cas II-gly on signaling and apoptosis. Effects of Cas II-gly on signaling and apoptosis. Cas II-gly has activating effects on MAPK and Ca^2^ + signaling, which in turn activate apoptosis. Inhibition of Wnt and TGF-*β* also lead to apoptosis induction. Mixed effects in other pathways also contribute to the perturbation of the apoptosis pathway. Additionally, perturbation of TGF-*β* may lead to other antitumorigenic actions of Cas II-gly, while the perturbation of Ca^2^ + signaling may lead to cardiotoxic side effects, which should be evaluated in a preclinical setting.

### Changes in pathways related to metabolism

Seven smaller communities, ranging from 3 to 8 pathways in size, were found to be integrated by pathways related to the biotransformation of a variety of compounds.

A community of five activated pathways was found, containing pathways related to drug and xenobiotic metabolism by cytochrome P450 (golden nodes with red borders at the bottom center of Metabolism frame in Fig. 1). The other perturbed pathways in this community are related to the metabolism of starch and sucrose, retinol, and porphyrin. The perturbation of these pathways by Casiopeina II-gly suggests their relative importance in the biotransformation of this compound. The cytochrome P450 (CYP) family plays an important role in the phase I biotransformation of xenobiotics [46, 48]. A role of cytochrome P450 in Casiopeina II-gly metabolism would have clinical implications related to bioavailability and disposition, as well as potential drug interactions, as it has been observed with other anticancer chemotherapeutics [44, 56]. Indeed, a recent work shows an inhibitory effect of a related compound of Casiopeina II-gly (Cas III Ea) on the activity of CYP1A1 [10].

There are few reports of porphyrin metabolism alterations following treatment with chemotherapeutic agents such as cisplatin [5], doxorubicin [39], docetaxel and trastuzumab [2]. As porphyrins are able to form complexes with metallic ions similar to Casiopeina II-gly, the use of this metabolic pathway for biotransformation would not be surprising. Meanwhile, retinoids have several roles in biological functions related to cancer [9], making them attractive pharmacological targets [47], while crosstalk between retinoid signaling and the induction of CYP exists [13]. As such, Casiopeina II-gly perturbation of this pathway may lead to interesting chemotherapeutic applications [17].

Carbohydrate metabolism was perturbed, as seen in the community composed by the upregulated pathways of glycolysis and gluconeogenesis, fructose and manose, galactose, pyruvate, and pentose phosphates, all of which were upregulated by Casiopeina II-gly. Another pathway in this community, glycan synthesis, was found to be downregulated. Glycans are known to be involved in several features that show deregulation in cancer cells [8, 40]. This is an example of a potential off-site target being modulated by Casiopeina II-gly.

Meanwhile, possible alterations to lipid metabolism were seen in a community composed by the PPAR signaling pathway (overexpressed) and the peroxisome pathway (underexpressed). Interestingly, in this community we also found the ABC transporters (overexpressed), which also play a role in drug disposition and possibly in particular features of cancer [18]. A related community, containing the linoleic and arachidonic acid pathways, was also upregulated.

Amino acid metabolism was found upregulated, as seen in the perturbation of the pathways of glycine, serine, and threonine, phenylalanine. The community containing these (light blue nodes at the bottom right of Fig. 1) also contained the nitrogen metabolism pathway, which also exhibited up-regulation. Another community contained the downregulated pathways of glycosaminoglycan synthesis and aminoacyl tRNA synthesis, and the pathways of sulfur metabolism and selenoamino metabolism exhibiting a mixed perturbation. Finally, a community of downregulated pathways related to purine, pyrimidine, RNA polymerase, and homologous recombination. Overall, the perturbation of these pathways gives a general idea of the global effects that Casiopeina II-gly by damaging DNA and how these affect cellular metabolism, not in an absolute fashion unlike cisplatin [55]. A more detailed metabolic profile of Casiopeina activity could lead to identify metabolic targets which may complement this line of treatment [61]. A simplified diagram of these relationships can be seen in Figure 3.

### Upregulation of processes related to immune response

Twelve other deregulated pathways formed their own community. Eight of these pathways describe disease conditions: asthma, allograft rejection and graft versus host disease, leishmaniasis, viral myocarditis, lupus, type I diabetes mellitus and autoimmune thyroid disease. The other four are antigen presentation, complement and coagulation cascades, cell adhesion molecules, and the hematopoietic cell lineage. The common element of these pathways is the involvement of immune response. With this in mind, the upregulation of these pathways by Casiopeina II-gly is indicative of the potential of this compound to elicit an immune response.

**Figure 3:**
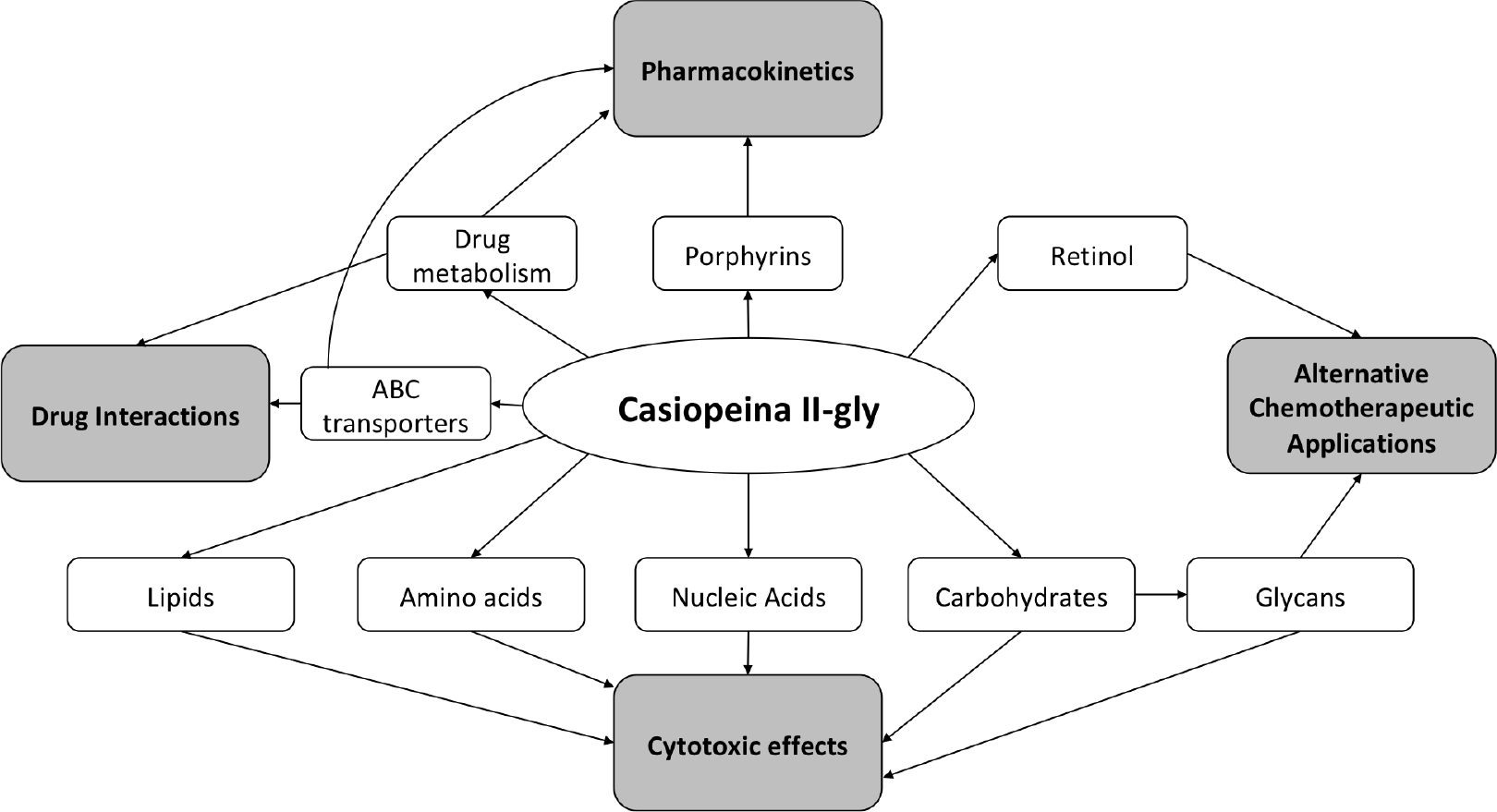
Effects of Cas II-gly on metabolic pathways. Cas II-gly directly affects metabolic pathways for different biomolecules, including aminoacids, lipids, carbohydrates, and nucleic acids, indicative of its cytotoxic effect. Cas II-gly also directly affects pathways related to drug metabolism, including the ABC transporters, which may be involved in the pharmacokinetics and possible drug interactions of Cas II-gly. Porphyrin metabolism is in turn affected by this drug, which may be related to metallic ions metabolism, also affecting their pharmacokinetics. Cas II-gly’s effects on retinol and glycan metabolism may indicate alternative chemotherapeutic applications for this compound.

The evasion of immune response is one of the hallmarks of cancer [23]. As such, the activation of an effective immune response is desirable in cancer treatment. Recently, the role of molecules belonging to the immune system as secondary targets through which known chemotherapeutic compounds exhert antitumor activity has been discussed [6]. The potential of Casiopeina II-gly to induce these complementary anticancer responses could have a positive impact in its clinical efficacy. A simplified diagram of these relationships can be seen in Figure 4.

**Figure 4:**
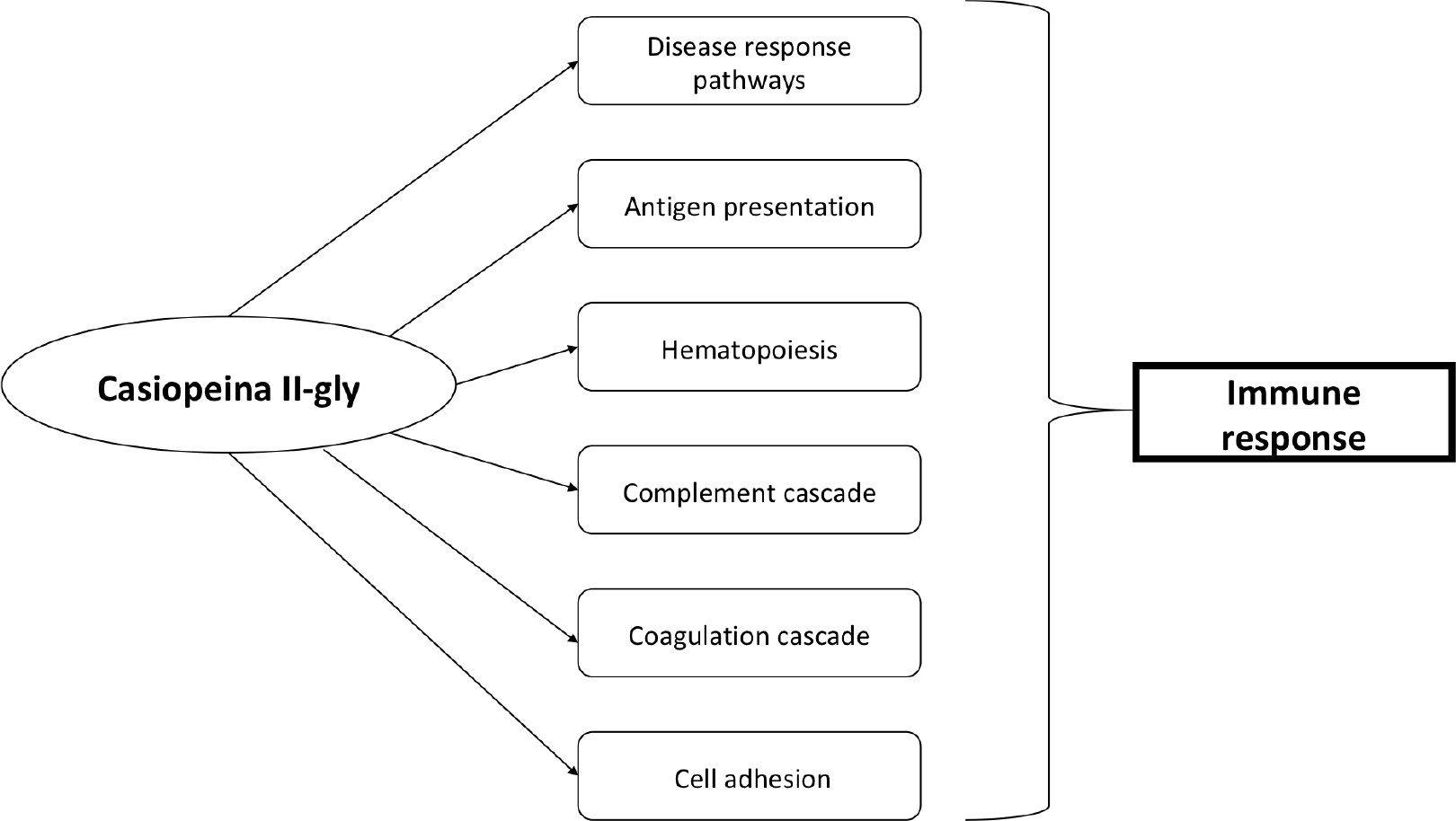
Effects of Cas II-gly on disease response. Cas II-gly has direct effects on pathways related to response to antigen presentation, hematopoiesis, complement and coagulation cascades, and cell adhesion. Cas II-gly also affects pathways associated with the response of multiple disease signatures. Taken together, these perturbations indicate a immune response action ellicited by Cas II-gly.

### Basal transcription and folate biosynthesis

Finally, two more pathways were identified to be perturbed by Casiopeina treatment. Unlike the other pathways, these have no shared molecules with any other pathway that exhibited any type of perturbation. In this sense, they are disconnected in the perturbed pathway network. First, we have downregulation of the basal transcription pathway. As we discussed above, the downregulation of transcriptional activity may be related to the general disruption of cell cycle that Casiopeina is exherting through DNA damage, reactive oxygen species, and mitochondrial dysfunction [43, 4, 29]. Action upon the folate biosynthesis pathway could be showing a compensatory mechanism after DNA damage, and open the possibility of benefit of therapy with antifolate chemotherapeutics such as metotrexate [49].

## Conclusions

By constructing and analyzing a network of pathways perturbed by Casiopeina II-gly *in vitro* treatment, we were able to identify relationships between these pathways that point to a potential for the compounds to exhibit effects beyond those previously described.

As expected from any cytotoxic agent, here we show that Casiopeina II-gly have effects on a wide range of biological processes, beyond its main mechanism of action. We show that many alterations in signaling pathways seem to be directly connected to the apoptosis induction mechanism of action.

We identified alterations to the expression of genes related to metabolic pathways involved in the processing of carbohydrates, lipids, nucleic acids, and aminoacids, as well as alterations to the metabolism of xenobiotics. Similarly, we identified deregulation of processes that point to the induction of an immune response, a response that can positively affect the cytotoxic efficacy of the compound *in vivo*.

Our model indicates that these three areas, namely, signaling/apoptosis, metabolism, and immunity exhibit interconnection and therefore can have effects upon each other. Moreover, our network model also identifies that these processes are organized in a modular structure, which opens the possibility for the development of combination therapies that integrate Casiopeina II-gly, to reduce adverse effects and increase efficacy.

We believe that by modeling the effects of Casiopeinas on gene expression at the pathway level, we may gain insights of functional effects of the compounds. It is worth to notice that this model does not rely on differentially expressed genes, which allows for a broader exploration of pathway deregulation, altough it limits its power to identify individual gene perturbations.

Our approach could be particularly useful to medicinal chemists, who can use this methodology to guide the evaluation of safety and efficacy of medicinal compounds. This model is not predictive, but rather, it serves as a hypothesis generation system which can lead to the design of new experiments, to gain a stronger understanding of the effects of a new compound.

